# Identification and structural characterization of small-molecule inhibitors of PINK1

**DOI:** 10.1101/2023.10.02.560389

**Authors:** Shafqat Rasool, Tara Shomali, Luc Truong, Nathalie Croteau, Simon Veyron, Jean-François Trempe

## Abstract

Mutations in PTEN-induced putative kinase 1 (PINK1) cause early-onset autosomal recessive Parkinson’s Disease (PD). PINK1 is a Ser/Thr protein kinase which functions as a mitochondrial damage sensor and initiates mitochondrial quality control by accumulating on the damaged organelle. There, it will autophosphorylate and then phosphorylate ubiquitin chains, which in turn will recruit and activate Parkin, and E3 ubiquitin ligase also implicated in PD. Ubiquitylation of mitochondrial proteins leads to the autophagic degradation of the damaged organelle. Pharmacological modulation of PINK1 constitutes an appealing avenue to study its physiological function and develop PD therapeutics. In this study, we used a thermal shift assay to identify small-molecule inhibitors of PINK1. In vitro kinase activity assays demonstrate that these molecules are ATP competitive inhibitors that block ubiquitin phosphorylation. PRT062607 (a SYK inhibitor) is the most potent inhibitor of PINK1 in our screen and has an *IC*_*50*_ of 2 μM against insect PINK1 and 1 μM in HeLa cells expressing human PINK1. The crystal structures of PINK1 in complex with PRT062607 or CYC116 reveal how the compounds interact with the ATP-binding pocket. PRT062607 notably engages with the catalytic aspartate (type-1 inhibition) and causes a destabilization of insert-2 at the autophosphorylation dimer interface. Our findings provide a scaffold for the development of more selective and potent inhibitors of PINK1 that can be used as chemical probes.

## Introduction

Mutations in PINK1 result in early onset autosomal recessive Parkinson’s Disease (PD) (1). PINK1 functions as a sensor of mitochondrial damage and works together with its effector, Parkin, to remove damaged components of the mitochondrial network (2). Upon mitochondrial damage such as depolarization, reactive oxygen species (ROS), or unfolded proteins, PINK1 accumulates on the outer mitochondrial membrane (OMM) and forms a supramolecular complex with the translocase of outer mitochondrial membrane (TOM) (3-7). Accumulation of PINK1 molecules results in its oligomerization and trans autophosphorylation at a conserved and unique serine (Ser228) in the N-lobe of the PINK1 kinase domain (8, 9). Autophosphorylation at this site results in the conformational rearrangement of the kinase domain of PINK1 and unlocks its ability to bind and phosphorylate ubiquitin (Ub) tethered to adjacent OMM proteins at Ser65 (8, 10-12). Phosphorylated Ub acts as a receptor for Parkin on the OMM (13-15), where it recruits Parkin. This in turns enables Parkin phosphorylation by PINK1 on its Ubl domain at Ser65 (16-18), which increases its E3 ligase activity leading to the ubiquitylation of OMM proteins and initiation of autophagy or formation of mitochondria derived vesicles (MDVs) carrying damaged cargo (19-21). In a physiological context, PINK1/Parkin do not mediate global basal mitophagy, but rather participate in the turnover of a subset of mitochondrial proteins upon stress (22-24). PINK1 suppresses activation of innate and adaptive immune responses that are triggered by mitochondrial stress or bacterial infection (25, 26). Hence, the activation of PINK1 is important to prevent accumulation of damaged mitochondrial components that can elicit deleterious inflammatory and autoimmune responses that lead to PD.

Pharmacological modulation of PINK1 presents an exciting avenue for a new PD therapeutic. Even in the absence of Parkin, PINK1 alone is sufficient to activate mitophagy, suggesting that PINK1 activation is a critical factor for mitochondrial quality control (19). Kinases are also a highly druggable class of protein presenting numerous orthosteric and allosteric binding sites that can accommodate small-molecules as therapeutic candidates (27). Inspired by the observation that kinetin triphosphate (KTP) can activate PINK1 *in vitro* and in cells (28), Mitokinin has developed a small molecule, MTK458, that sensitizes PINK1 to lower levels of mitochondrial damage and activates PINK1/Parkin-mediated mitophagy in cells (29). Additionally, MTK458 also mitigates lipopolysaccharides (LPS)-induced inflammation and propagation of α-synuclein fibrils in mice, demonstrating the potential of PINK1 activators as therapeutic candidates for PD. However, the molecular mechanism by which MTK458 activates PINK1 is not understood.

Biochemical and cellular studies on PINK1 have entirely relied upon the use of genetic tools (knockdowns and knockouts) and mitochondrial damaging system to study the function of PINK1 (30). Mechanistic dissection of PINK1 function demands more sophisticated tools that can delineate PINK1 expression from mechanistic phenomenon such as stabilization on mitochondria, activation of kinase activity and binding to substrate. In the study of kinase function, specific inhibitors of kinases are often employed to study their role in the cell. However, in the absence of specific small-molecule binders of PINK1, it is not possible to study the function of PINK1 in this manner. There is thus a need to develop tool compounds against PINK1 to allow for acute inhibition of its active in vivo. Tool compounds are also useful for structural studies and in some cases can serve as scaffold for the development of activators such as paradoxical agonists (31).

Recently published structures of insect PINK1 have shed light on the mechanism of PINK1 activation by autophosphorylation via a novel dimerization interface that enables autophosphorylation at Ser228 in the N-lobe (8, 32). These structures have also identified PINK1-specific regions that are essential for PINK1 activation such as insert-2 for autophosphorylation or insert-3 for Ub/Ubl substrate recognition. These studies with insect variants such as *Tribolium castaneum* PINK1 (TcPINK1) have also helped unlock protein engineering and methodological approaches to produce stable recombinant protein, which is critical for screening small-molecule binders and study their structures by cryo-electron microscopy (cryoEM) or crystallography.

In this study we report the screening and discovery of small-molecule binders of insect and human PINK1. Using thermal shift assays, we screened a library of kinase inhibitors and identified candidates targeting TcPINK1. The most potent compound, PRT062607, inhibits TcPINK1 with *in vitro IC*_*50*_’s in the 1-2 μM range. The crystal structure of TcPINK1 bound to PRT062607 reveals that it binds in the ATP binding site and captures the kinase in an activated state, forming interactions with the hinge region as well the activation loop. *In organello* assays with isolated damaged mitochondria as well as cell-based assays showed that PRT062607 and related compounds also inhibit human PINK1 and reduce its ability to phosphorylate Ub. These small molecules can serve as scaffolds to produce highly potent and specific inhibitors of PINK1 to be employed as tool compounds.

## Results

### Identification of PRT062607 as an inhibitor of TcPINK1 ubiquitin kinase activity

Our goal is to identify a ligand that can be used as a starting point for the development of a chemical probe for human PINK1. The ATP-binding site of kinases is well conserved and thus we hypothesized that screening a library of kinase inhibitors would enable identification of one or more inhibitors that inhibit PINK1. Binding of inhibitors to kinases typically lead to their stabilization, via interaction with elements of the active site. We used a thermal shift assay to screen for ligands that bind PINK1, an approach that has been used previously to identify kinase ligands (33). The kinase domain from human PINK1 cannot be purified in its active form from bacteria. We thus used TcPINK1, which can be purified in its active form following expression in bacteria and has 48% sequence identity with human PINK1 in the kinase domain (10, 34). Using the fluorescent dye Sypro-orange, we find that the melting temperature (*T*_*m*_) of TcPINK1 is 45.1 °C (Suppl. Fig. 1). We then used this assay to screen over 400 compounds at 100 μM and identified 8 compounds that increased the melting temperature (*T*_*m*_) by more than 1.5 °C (Fig. 1A). These compounds include the pan-kinase inhibitor staurosporine, which induced stabilization by 4.8 °C. To determine the relative potency of these compounds towards inhibiting TcPINK1, we conducted an *in vitro* kinase assay using tetraubiquitin (Ub_4_) as a substrate in the presence of 100 μM compound (Fig. 1B). Quantification shows that PRT062607 inhibits Ub_4_ phosphorylation most strongly. We confirmed that PRT062607 inhibits TcPINK1 by performing a similar kinase assay using monoubiquitin and assessing product formation using Phos-tag gels (Fig. 1C). PRT062607, also known as BIIB057 or P505-15, is a potent inhibitor (*IC*_*50*_ = 1 nM) of the SYK tyrosine kinase (35). We thus tested additional SYK inhibitors that are structurally related to PRT062607, namely PRT060318 and TAK-659, as well as a racemic mixture of the “core” of PRT062607 (Fig. 1D). We found that they all inhibit the kinase activity of TcPINK1 (Fig. 1B).

**Figure 1.**
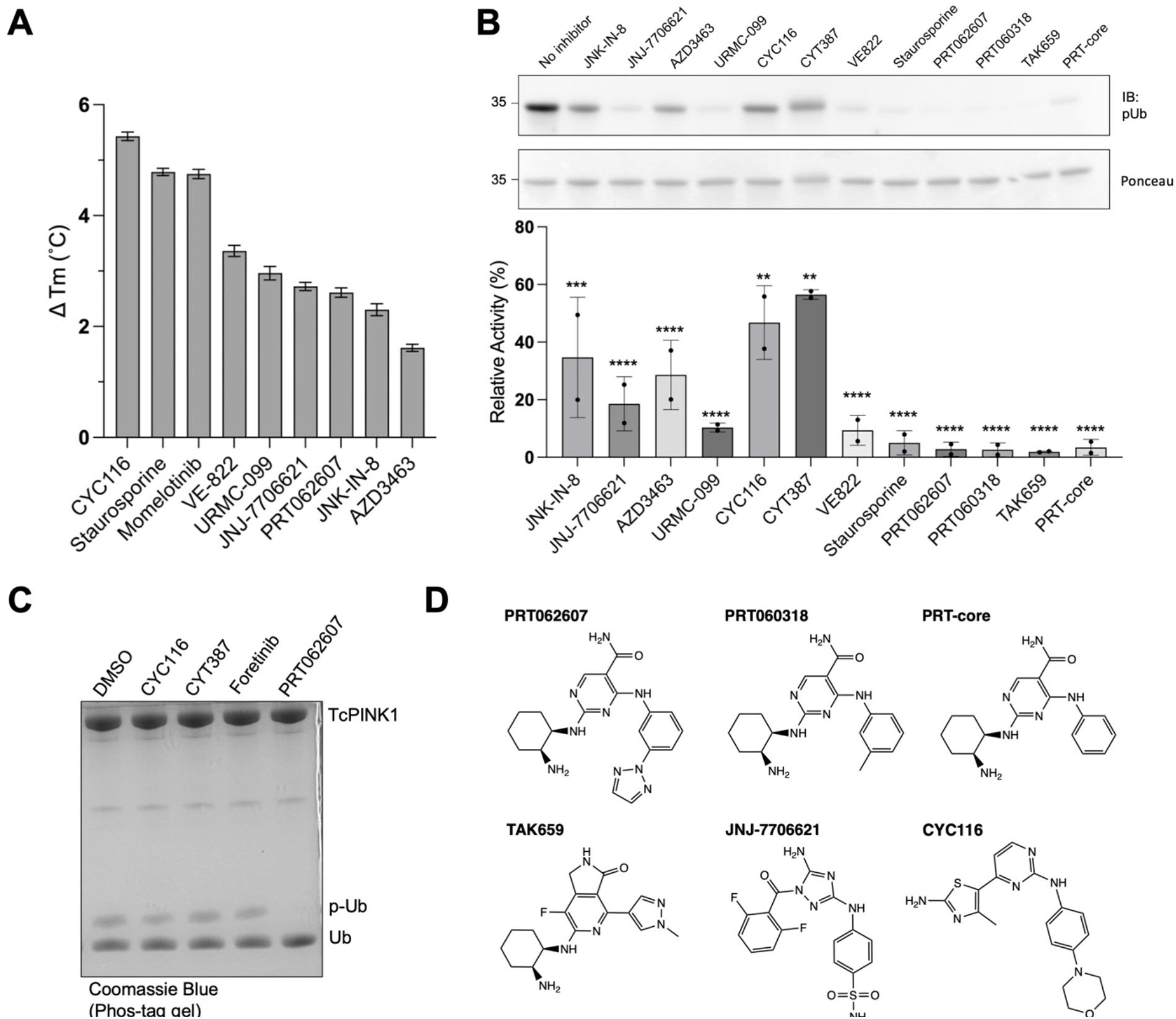
Identification of PRT062607 as an inhibitor of PINK1 ubiquitin kinase. **(A)** Thermal shift assay screen of kinase inhibitors against *Tribolium castaneum* PINK1 (TcPINK1) with melting temperature increase of 1.5°C and above. **(B)** In vitro ubiquitin kinase assays with TcPINK1 and tetraubiquitin (Ub_4_). Reactions were analyzed by immunoblotting against phospho-ubiquitin (pUb). Formation of pUb is significantly impaired at alpha = 0.05. ns at P > 0.05, * at P ≤ 0.05, ** at P ≤ 0.01, *** at P ≤ 0.001 and **** at P ≤ 0.0001. The ponceau staining of the above gel is shown as loading control. **(C)** In vitro ubiquitin kinase assays with 5 μM GST-TcPINK1 and 30 μM ubiquitin, as well as 1 mM ATP at 37°C. Reactions were analyzed using a phos-tag gel and stained by Coomassie blue. **(D)** Chemical structures of identified PINK1 inhibitors.

### PRT062607 is an ATP-competitive inhibitor of PINK1

To determine the mechanism of action of PRT062607 and related compounds, we incubated ^13^C-labeled adenosine triphosphate (ATP) with recombinant TcPINK1 and monitored hydrolysis by acquiring ^13^C-^1^H HSQC NMR spectra. TcPINK1 displays ATP hydrolysis activity even in the absence of a ubiquitin substrate, as observed by the appearance of a peak corresponding to ADP (Fig. 2A). Addition of excess PRT062607 prevented ATP hydrolysis, consistent with PRT062607 competing with ATP. Furthermore, we found that PRT062607 also inhibited TcPINK1 autophosphorylation, which would also be expected from an ATP-competitive inhibitor (Fig. 2B). Given that the inhibitors compete with ATP, we thus sought to determine the potency of the various inhibitors using the Kinase Glo assay, which measures consumption of ATP using luminescence for increased throughput. To determine the optimal conditions for *IC*_*50*_ determination, we first conducted an assay in the presence of different concentrations of TcPINK1 and fixed ATP concentration (Suppl. Fig. S2A). We found an *EC*_*50*_ around 1 μM of TcPINK1, which we have used for all the inhibition assays. TcPINK1 binds to a non-hydrolyzable ATP analog (AMP-PNP) with a *K*_*d*_ of 96 μM (Fig. 2C), which is within the normal range of *K*_*d*_ or *K*_*m*_ values for active Ser/Thr kinases and ATP (36). We thus used a low concentration of ATP of 10 μM to characterize ATP competitive inhibitors, in which case the *IC*_*50*_ should approximately be equal to the *K*_*i*_ constant (*IC*_*50*_ = *K*_*i*_ (1+[ATP]/*K*_*m*_). In the absence of inhibitors, both unphosphorylated and monophosphorylated TcPINK1 hydrolyze ATP to similar extent in 5 min (Suppl. Fig. S2B). PRT062607 was then titrated against both forms and was found to inhibit ATP hydrolysis with an *IC*_*50*_ around 2 μM in both cases (Fig. 2D), implying that the phosphorylation status of PINK1 does not affect inhibitor binding. The closely related compound PRT060318, with a methyl group replacing the triazole, likewise inhibited TcPINK1 with an *IC*_*50*_ of 2.5 μM. The other inhibitors (TAK-659, JNJ-7706621 and CYC116) had lower potencies, with *IC*_*50*_ between 6 and 65 μM. PRT02607 is thus the most potent inhibitor of TcPINK1 amongst the hits identified in the thermal shift screen.

**Figure 2.**
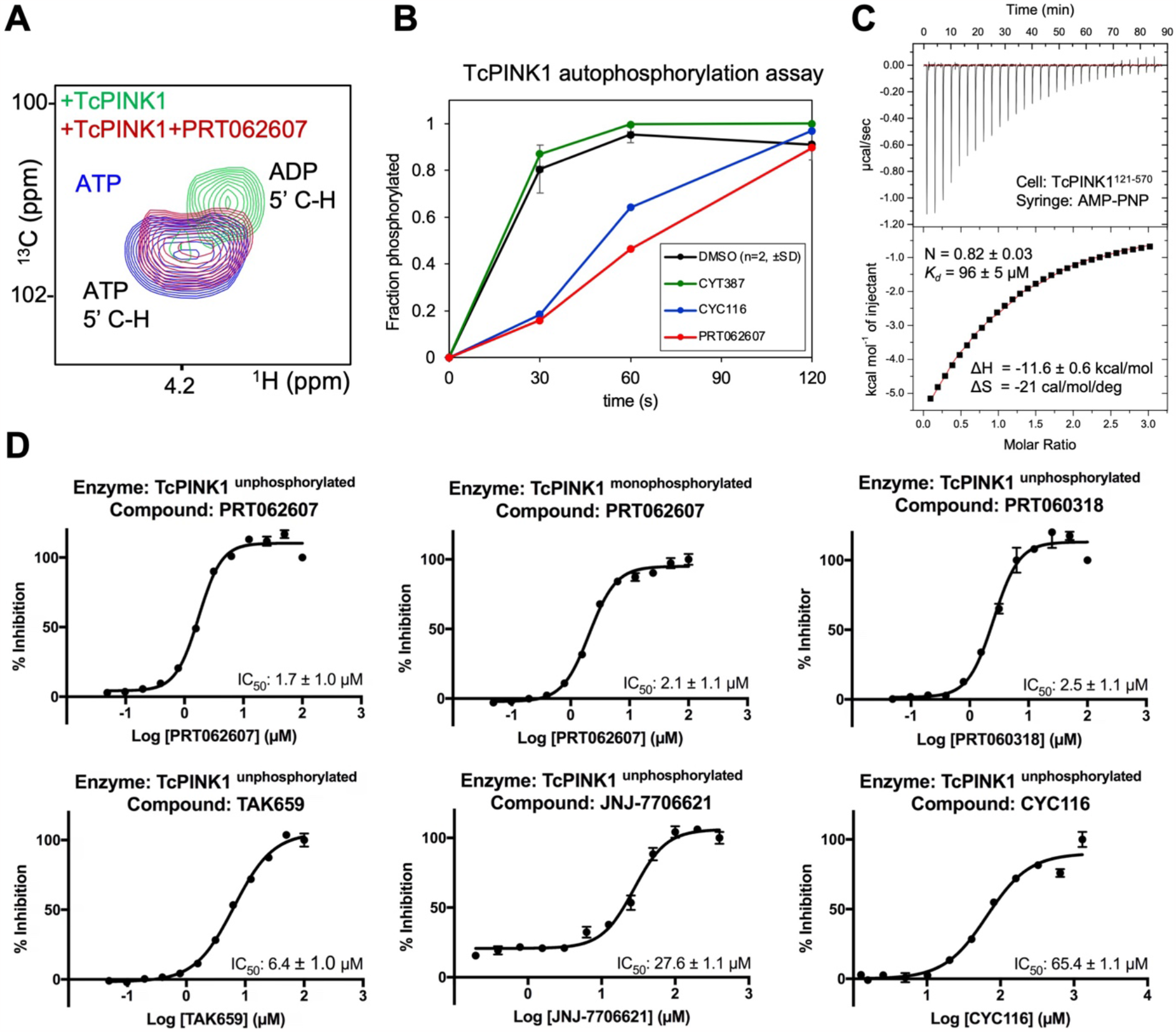
PRT062607 and related compounds are competitive inhibitors of ATP hydrolysis. **(A)** ^13^C-^1^H HSQC NMR of ^13^C-ATP (100 μM), showing the cross-peak of 5’ C-H. Incubation for 1h with 1 μM TcPINK1 leads to complete hydrolysis to ADP, which is inhibited by 100 μM PRT062607. **(B)** Inhibition of TcPINK1 autophosphorylation (5 μM), in the presence of 100 μM different compounds and 1 mM ATP at 30°C. The phosphorylated fraction was determined by LC-MS. **(C)** Isothermal calorimetry (ITC) of TcPINK1 (cell) titrated with AMP-PNP (syringe). Data were fitted to a one-site model, with a Chi^2^/DoF = 1612. **(D)** *IC*_*50*_ determination for different inhibitors of TcPINK1 using the Kinase Glo luminescence assay. Measurements were taken 5 minutes after TcPINK1 was incubated with different concentrations of the inhibitors and 100 μM ATP. The error bar indicates the standard deviation (n=3). The results were normalized and are presented as percent inhibition of TcPINK1 compared to no inhibitor control.

### PRT062607 inhibits human PINK1 *in organello* and in cells

To determine whether the compounds we identified against TcPINK1 also inhibit human PINK1, we performed a Ub_4_ kinase assays with isolated mitochondria from human HeLa cells that were treated with carbonyl cyanide m-chlorophenyl hydrazone (CCCP), a protonophore that induces PINK1 accumulation on mitochondria. The results show that PRT062607 and PRT060318 are the most potent inhibitors of human PINK1 kinase activity (Fig. 3A). To determine the potency of PRT062607 inhibiting human PINK1 in cells, we treated HeLa S3 cells overexpressing PINK1-3HA with CCCP for 4h in the presence of different PRT062607 concentrations, followed by immunoblotting against pUb and HA (Fig. 3B). CCCP induces formation of a smear on the pUb blot, characteristic of multiple pUb being conjugated to mitochondrial outer membrane proteins. PRT062607 inhibits ubiquitin phosphorylation in cells in a concentration-dependent manner, with an *IC*_*50*_ of 1.2 μM (95% CI: 0.5-3.4 μM). This is consistent with the known membrane permeability of PRT062607, which can diffuse and inhibit the cytosolic kinase domain of PINK1 bound to mitochondria. We also observe that the levels of PINK1 are not substantially affected by PRT062607, implying that the observed reduction in pUb is not caused by a reduction in PINK1 levels.

**Figure 3.**
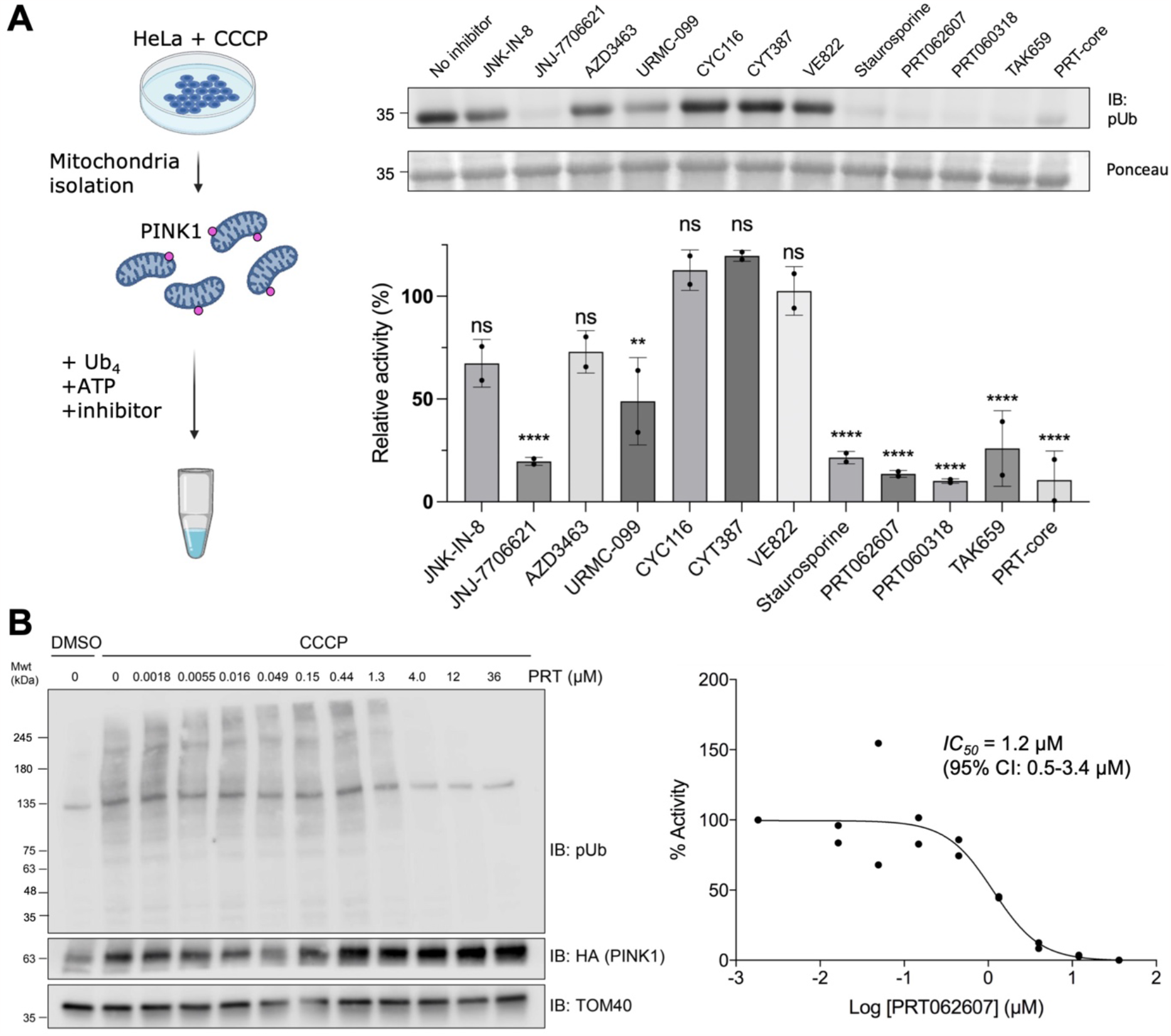
PRT062607 and related compounds inhibit human PINK1 *in organello* and in cells. **(A)** In organello ubiquitin kinase assays with accumulated HsPINK1 and tetraubiquitin (Ub_4_). Reactions were analyzed by immunoblotting against phospho-ubiquitin (pUb). Formation of pUb is significantly impaired at alpha = 0.05. ns at P > 0.05, * at P ≤ 0.05, ** at P ≤ 0.01, *** at P ≤ 0.001 and **** at P ≤ 0.0001. The ponceau staining of the above gel is shown as loading control. **(B)** *IC*_*50*_ generated from HeLa S3 cells transfected with endogenous HA-tagged HsPINK1 treated for 4 hours with 20 μM CCCP. Reaction was performed twice as biological replicates. Reactions were analyzed by immunoblotting against phospho-ubiquitin (pUb). The 100% activity baseline was measured as no PRT062607 with CCCP treatment. Immunoblots against blots HA and TOM40 are shown as loading controls.

### Crystal structure of PINK1 bound to small-molecule inhibitors

In order to obtain a detailed understanding of how the inhibitors identified above bind to PINK1, we co-crystallized the cytosolic domain of TcPINK1 (121-570) using the crystallization construct we previously used to obtain the structure of TcPINK1 in the apo and AMP-PN bound forms (8). This construct was generated by mutating 6 non-conserved surface exposed aromatic residues to alanine to increase solubility. While we managed to obtain crystals for TcPINK1 in presence of PRT062607, PRT060318, CYC116 and TAK-659, only the co-crystals with PRT062607 yielded diffraction data at 2.9 Å resolution that could be used to obtain a structure (Table 1). Introduction of two additional solubilization substitutions to non-conserved hydrophobic surface residues allowed us to determine the structure of TcPINK1 in complex with CYC116 at 3 Å resolution (Table 1). Ligand omit maps for both inhibitors show electron density for nearly all atoms, except the most distal elements exposed to the solvent (Suppl. Fig. 3A).

**Table 1.** X-ray crystallography data collection and refinement statistics.

|  | **TcPINK1<sup>121-570</sup>: PRT062607**<br>**(PDB: 8UCT)** | **TcPINK1<sup>121-570</sup>: CYC116**<br>**(PDB: 8UDC)** |
| --- | --- | --- |
| Wavelength (Å) | 0.987 | 0.987 |
| Resolution range (Å) | 90.93–2.93 (2.98–2.93) | 54.60–2.99 (3.16–2.98) |
| Space group | P6<sub>1</sub>22 | P6<sub>1</sub>22 |
| Unit cell dimensions (Å) | 53.216 53.216 545.580 | 52.99 52.99 545.97 |
| Unit cell angles (deg) | 90.0 90.0 120.0 | 90.0 90.0 120.0 |
| Total reflections | 196,146 (8,782) | 184,476 (30,457) |
| Unique reflections | 11,014 (521) | 10600 (1611) |
| Multiplicity | 17.8 (16.9) | 17.4 (18.9) |
| Completeness (%) | 100 (100) | 100 (100) |
| Mean I/sigma(I) | 6.1 (0.5) | 3.5 (0.6) |
| Wilson B-factor | 60 | 71.3 |
| R-merge | 0.605 (8.848) | 0.613 (9.276) |
| CC<sub>1/2</sub> | 0.995 (0.551) | 0.991 (0.325) |
| Reflections used for R-free | 546 | 514 |
| R-work | 0.2834 | 0.2713 |
| R-free | 0.3052 | 0.3221 |
| # non-hydrogen atoms | 3051 | 3046 |
| Macromolecules | 2981 | 3016 |
| Ligands | 47 | 26 |
| Water | 23 | 4 |
| Protein residues | 391 | 397 |
| RMS (bonds, Å) | 0.007 | 0.011 |
| RMS (angles, deg) | 0.776 | 1.221 |
| Ramachandran favored (%) | 94 | 90 |
| Ramachandran allowed (%) | 6 | 10 |
| Ramachandran outliers | 0 | 0 |
| Clashscore | 5.50 | 8.5 |
| Average B-factor | 77.0 | 93.0 |
| Macromolecules | 71.0 | 76.0 |
| Ligands | 129.5 | 169 |
| Solvent | 79.6 | 85.0 |

The overall architecture of TcPINK1 bound to these inhibitors is identical to the previously published insect PINK1 structures with a folded bi-lobular kinase domain followed by a helical C-term extension (CTE) (8, 10-12, 32, 37). As expected from our kinase assays, both inhibitors occupy the ATP-binding site of PINK1 between the two lobes. Similar to previous structures, the inhibitor bound structures feature an active kinase domain with all of its structural hallmarks; structured activation loop (A-loop) with the ‘DFG-in’ conformation, folded αC helix adjacent to the active site with its Asp217 forming a salt bridge with Lys196 in β3 sheet, intact R-spine formed by hydrophobic contacts between the buried side chains of His335 in the catalytic HRD motif, Phe360 in the DFG motif and Val218 in the αC helix, and C-spines formed by hydrophobic contacts between the side of Leu344 in the C-lobe, the aromatic scaffold ring in both inhibitors and Val176 in β2 (Fig. 4A). These observations confirm that both molecules act as type-I inhibitors of PINK1.

**Figure 4.**
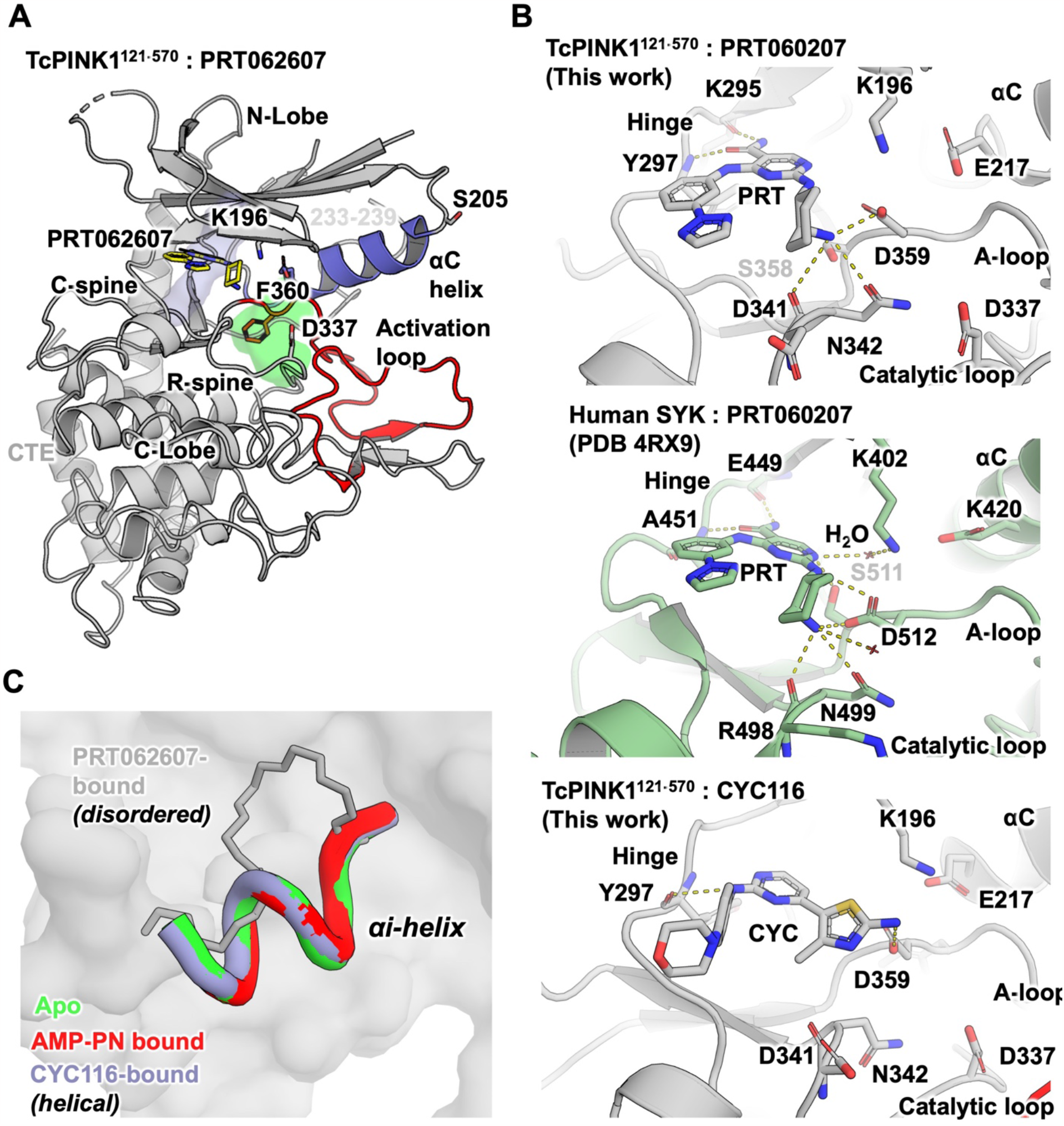
Crystal structures of TcPINK1 with PRT062607 and CYC116 reveal basis for inhibition. **(A)** Crystal structure of TcPINK1^121-570^ (cartoon) in complex with PRT062607 (yellow sticks). Structure of TcPINK1 bound to PRT060207 showing ordered activation loop (red) and αC helix (slate), intact R-spine (green) and C-spine (blue). The insert-2 segment spanning a.a. 233-239 is disordered. **(B)** Active site of TcPINK1 bound to PRT060207 (top) and CYC116 (bottom) showing their interactions with the hinge, A-loop and catalytic loop of each protein. The active site of SYK bound to PRT060207 (PDB 4RX9, middle) show similar interactions with the hinge, A-loop and catalytic loop. Dashed lines indicate H-bonds. **(C)** Change in conformation observed in the αi helix upon binding PRT062607. Superposition of the structures of TcPINK1^121-570^ in the apo form (PDB 7MP8, green), or bound to AMP-PN (PDB 7MP9, red), PRT062607 (this work, grey), or CYC116 (this work, violet).

Both inhibitors form polar interactions with the hinge and active site region of the kinase (Fig. 4B, top and middle). In PRT060207, the carboxy acetamide group forms H-bonds with the backbone carbonyl and amide groups of Lys295 and Tyr297 respectively in the hinge region. The amine group of the 2-aminocylohexane group forms multiple H-bonds with surrounding residues including the side chains of Ser358 (DFG-1) and Asp359 (DFG motif) in the activation loop, as well as Asn342 and the backbone carbonyl of Asp341 downstream of the catalytic HRD motif (335-337). These interactions are identical to the interactions of PRT060207 with SYK (Fig. 4 B, bottom; PDB 4RX9) (38). On the other hand, CYC116 interacts with the backbone carbonyl of Tyr297 in the hinge with the amine group connecting the pyrimidine ring and the phenyl group. H-bonds with Asp359 are formed by the amino group on the 5-membered 2-amino-4-methyl-1,3-thiazol-5-yl ring moiety. However, the latter group is not charged, which may explain the lower potency of CYC116.

Crystallographic symmetry analysis of PRT062607 and CYC116 bound structures reveal the same dimeric face-to-face trans autophosphorylation complex seen in our previous structure with the Ser205 from each protomer extending to the active site of the opposing promoter and interacting with the Asp337 (Suppl. Fig. S3B). This dimer interface is notably supported by distal interactions between the αi helix in Insert-2 and a.a. 378-397 in the activation segment, including salt bridges between Glu379 from one protomer and Arg216 and Arg241 of the other protomer (8). Interestingly, while CYC116 maintains a helical αi helix identical to the one observed in our previous structures, this segment (a.a. 233-239) is disordered and retracted in the structure with PRT062607, rendering it unable to interact with the activation segment of the opposing protomer (Fig. 4C). This conformational change induced by the drug would thus impair formation of the *trans* autophosphorylation dimer.

To understand the binding mechanism of TAK-659 and JNJ-7706621, we generated docking models of TcPINK1 with these compounds (Suppl. Fig. S3C, left and right). Similar to the structure with PRT060207, TAK-659 forms H-bonds with the backbone of Lys295 and Tyr297 via the polar groups in its pyrrolo group and central fluoro-pyridine ring. The aminocyclohexane ring forms the same interactions with the activation and catalytic loops. JNJ-7706621 forms three H-bonds with the hinge through its central amino-triazole group, instead of two in the case of PRT062607-related compounds. However, owing to a lack of polar groups on the difluorophenyl ring, no interactions are observed with the activation or catalytic loops. Interestingly, the polar sulfonamide group is within interacting distance of Asn300 upstream of the hinge. These additional H-bonds with the hinge may explain why JNJ-7706621 maintains potency in the absence of interactions with the DFG motif.

## Discussion

In this study, we identified several small-molecule inhibitors active against human PINK1 using a thermal shift assay directed at an insect ortholog. This is consistent with the high degree of sequence and functional conservation in PINK1 across metazoans. This highlights the usefulness of TcPINK1 as a model for the human enzyme. Yet, we observe differences between the patterns of inhibition in TcPINK1 and HsPINK1 (compare Fig. 1B and 3A). Notably, VE822 inhibits TcPINK1 (8% activity at 100 μM compared to control) but does not significantly impact HsPINK1. As we seek to develop more selective compounds towards HsPINK1, it will be important to establish a system for the purification of its active form for structural studies. Several groups have reported purification of the human enzyme from bacteria, yeast or insect cells (28, 34, 39, 40). Recently, a preprint from the group of Miratul Muqit reported that genetically-encoded incorporation of phosphoserine at position 228 increased the activity of recombinant HsPINK1 purified from bacteria (41). Yet, assays performed with recombinant HsPINK1 typically require radioactive ATP to detect ubiquitin kinase activity, since only a small fraction of the pool of enzyme is active, which would not be conducive to structural studies. Until a more robust recombinant human PINK1 construct is developed, the *in organello* Ub_4_ phosphorylation and cell-based assays we have developed here will remain useful tools to at least assess the potency of compounds towards HsPINK1.

To better characterize the inhibitors, we measured their *IC*_*50*_*’s* with the Kinase Glo luminescence-based assay. To our surprise, TcPINK1 was able to hydrolyze ATP in the absence of a substrate. This cannot be explained by autophosphorylation alone, since 1 μM of TcPINK1 phosphorylated at Ser205 was able to consume 9 μM of ATP, similarly to the unphosphorylated enzyme (Supplemental Fig. S2). This futile ATP hydrolysis must arise from a water molecule acting as a substrate and mediating a nucleophilic attack on the γ phosphate of ATP. Furthermore, during the optimization of the assay, we observed that 2 μM of TcPINK1 consumed 60% (300 μM) out of a 500 μM pool of ATP in 5 min. This implies that the turnover rate for futile ATP hydrolysis is at least 30 min^-1^, which is in the same range as the *k*_*cat*_ of 43.5 min^-1^ measured for autophosphorylation at Ser205 (8). This reflects the fact that PINK1 is always in an active DFG-in conformation and requires accumulation on the TOM complex to phosphorylate its substrate, rather than its active site changing conformation via phosphorylation or substrate binding, as is the case for most protein kinases.

Two compounds, PRT062607 and CYC116, were co-crystallized with TcPINK1. The structures confirmed that the inhibitors occupied the ATP-binding site of PINK1, affirming their role as Type-I inhibitors. The structures provide insights into the specific polar interactions formed with the kinase hinge and active site residues such as the DFG motif, which are similar but not identical to the interactions of these inhibitors with their cognate kinases. Comparatively, the SYK:PRT062607 complex also contains ordered water molecules, one of which mediates the interaction between the amino group connecting the central pyrimidine ring to the 2-aminocyclohexane and the active site Lys402 (Lys196 in TcPINK1) (Fig. 4B). We could not confidently identify electron density for water molecules in this region, which could be attributed to the lower resolution of our structure (2.9 Å compared to 1.75 Å for SYK:PRT062607). Alternatively, perhaps these interactions mediated through water molecules do not occur in PINK1, which could account for the lower potency of PRT060207 for PINK1 compared to SYK. Our structure also reveals potential interactions that a chemical probe could make to enhance selectivity against PINK1. For instance, the active site of TcPINK1 is lined with Asp341 (Asp366 in human), which is an arginine, lysine or alanine at the same position in tyrosine kinases including SYK (Arg498), FGR (Ala386), and FLTK3 (Arg815), or Ser272 in MLK1 (42). Addition of a positively charged group on the PRT scaffold to interact with Asp366 would enhance selectivity for PINK1. Future work will exploit these interactions to design selective PINK1 chemical probes.

It remains to be understood how PRT062607 binding to the active site results in a distal destabilization of the αi helix (Fig. 4C). Since the protein construct, buffer, crystallization condition and crystal form of the structure with PRT062607 are identical to the previous structures (apo or with AMP-PNP), this change in conformation is unlikely to originate from a packing artefact and likely represents a genuine allosteric conformational change induced by the drug, which would further contribute to inhibiting the enzyme. The disorder of the αi helix may explain why despite being a better inhibitor of PINK1 than CYC116, PRT062607 induces a smaller increase in the melting temperature of TcPINK1 in thermal shift assays. The αC helix is located between the ATP binding site and αi helix and thus could mediate the allosteric, perhaps via interactions with Lys196. This feature could be important to achieve PINK1 inhibition via both orthosteric ATP displacement and allosteric destabilization.

The PINK1 inhibitors we have discovered were all designed to target other kinases and have primarily been tested in cancer, vascular, and autoimmune diseases. Given the evolutionary pressure to preserve the catalytic function of tyrosine kinases and serine/threonine kinases such as PINK1, there are similarities in the fine structure of the ATP binding site which leads to promiscuity in binding (43). For instance, PRT062607 and PRT060318 were originally discovered as SYK inhibitors (35, 44). SYK is known for its role in B cell lymphoma mediated signalling (45). Despite its relatively high potency for PINK1, PRT062607 is not specific for PINK1, with *IC*_*50*_’s in the nanomolar range for the tyrosine kinases SYK (1 nM) and FGR (81 nM), as well as the Ser/Thr kinase MLK1 (88 nM). PRT060318 is also a potent inhibitor of SYK at 4 nM (44). Both inhibitors prevent thrombocytopenia and thrombosis in transgenic mouse models. TAK-659 also inhibits SYK and impairs CLL proliferation after B-cell receptor activation, similarly to the PRT compounds (46). TAK-659 reversibly inhibits SYK at 3 nM and FMS-like tyrosine kinase 3 (FLTK3) at 5 nM (47). CYC116 inhibits the Aurora Ser/Thr kinases (A, B and C) and the vascular endothelial growth factor receptor 2 (VEGFR2) tyrosine kinase and has been shown to delay tumour growth in vivo mice model (48). CYC116 can inhibit PINK1, but not as potently as the PRT compounds, making CYC116 an unlikely scaffold for developing a selective PINK1 probe. Finally, JNJ-7706621 is a dual inhibitor of the Ser/Thr cyclin dependent kinases (CDK1 and CDK2) and Aurora kinases, with *IC*_*50*_’s in the 3-250 nM range (49). In human HeLa cells, JNJ-7706621 was able to inhibit cell growth and proliferation and at high concentration, induces cytotoxicity. JNJ-7706621 is an interesting scaffold as it forms three H-bonds with the hinge, which would increase the residence time of the drug. As we seek to develop more potent PINK1 inhibitors based on these scaffolds, the challenge will be to reduce the potency against other kinases to achieve sufficient selectivity for their use as chemical probes.

Overall, the discovery of PRT062607 and other ATP-competitive inhibitors and their promising inhibitory activity against human PINK1 lay a strong foundation for future studies focused on designing selective chemical probes. These probes could be used to assess the involvement of PINK1 in specific biological responses and to dissect the mechanism of action of activator drugs targeting the PINK1/Parkin pathway.

## Experimental procedures

### Protein purification

#### Plasmids

The TcPINK1 plasmids used spans residues 121-570 or 128-570 and are codon optimized for expression in E.coli (10).TcPINK1 (121-570) ampicillin-resistant plasmids were co-expressed in BL21 DE3 E. coli with lambda phosphatase spectinomycin-resistant plasmid (Addgene plasmid #79748). We used either wild-type, kinase-dead D337N (10), or solubilizing constructs with 4, 6 or 8 mutations, as indicated for each experiment below. The plasmid of tetraubiquitin (Ub4) was ordered from GenScript. This plasmid contains the ubiquitin gene repeated four times and contains a N-term GST tag as well as a 3C HRV cleavage site.

#### Bacterial growth and induction of expression

A single colony of each co-transformation were used to inoculate a 10 mL starter culture of Lysogeny Broth (LB) with 0.1 mg/mL ampicillin and 0.05 mg/mL of spectinomycin; the cultures were incubated at 37°C overnight, shaking at 180 rpm. The starter culture was then added to 1 L of LB or M9 containing ^15^NH_4_Cl, both supplemented with 0.1 mg/mL ampicillin and 0.05 mg/mL spectinomycin and left to grow at 37°C, shaking at 180 rpm until the optical density at a wavelength of 600 nm (OD600) reached 0.8-1.0. Cultures were cooled down to 16°C and 300 μM of IPTG and 25 μM of MnCl_2_ were added. The cultures were then incubated at 16°C overnight, shaking at 180 rpm. Cells were harvested by centrifugation at 3500 rpm for 30 minutes at 4°C. The resulting pellets were resuspended in 30 mL of lysis buffer consisting of 0.025 mg/mL DNAse, 5 mM MgCl_2_, 1 mM PMSF, 0.1 mg/mL lysozyme, 0.2% Tween 20, 10% glycerol, in 300 mM NaCl and 50 mM Tris-HCl pH 8. The resuspended pellets were sonicated at 4°C for 20 seconds on-off intervals repeated 8 times.

#### Cell lysis and protein purification

After cell lysis and sonication, the bacteria were spun down at 15000 rpm for 20 minutes at 4°C to separate the cell debris and insoluble protein from the soluble fractions. The clarified cell lysate (supernatant) was then incubated for 1 hour at 4°C on a rotating platform with 2 mL Sepharose 2B resin (GE Healthcare Life Sciences) suspended in 2 mL kinase buffer (50 mM Tris, 300 mM NaCl, 3 mM DTT, pH 8.0). After washing the resin repeatedly with kinase buffer (wash fractions), the protein of interest was eluted using 20 mM glutathione (Bio Basics Canada) and concentrated by centrifugation using 15 kDa Centrifugal Filter Units (Millipore). The resulting protein concentration was measured using the Denovix DS-11 spectrophotometer at 280 nm wavelength. The eluted protein was incubated with a 1:50 ratio of HRV-3C protease to protein at 4°C overnight to cleave the GST tag off the protein.

#### GST tag cleavage and size exclusion chromatography

The cleaved protein was injected on a Äkta pure (Cytiva) with the S75 HiLoad 16/600 Superdex column (Cytiva) and a 5 mL GSTrap attached at 0.7 mL/min equilibrated in the kinase buffer to separate GST from TcPINK1. The fractions eluted off the Äkta were further concentrated with the 15 kDa Centrifugal Filter Units (Millipore) according to the manufacturer’s instructions. The mass and phosphorylation status of each protein was verified using intact mass spectrometry.

### Thermal shift assay

TcPINK1^128-570^ D337N (kinase dead TcPINK1, or “TcPINK1 KD”) was used for the Thermal Shift Assay screening. Each of the 430 compounds from a SelleckChem Kinase Inhibitor Library (catalog #L1200) were combined at 100 μM in wells of a polypropylene 96-Well Tube Plates (Agilent) with TcPINK1 KD at 0.5 mg/mL (approximately 10 μM) and SYPRO Orange Protein Gel Stain 5,000X (Thermo Fisher Scientific) at 6X in a 300 mM NaCl, 20 mM Tris-HCl pH 8, 5 mM DTT, 5% DMSO buffer. The reaction mixtures were heated from 10°C to 70°C for approximately 1 hr by a QuantStudio 7 Pro Real-Time PCR System (Thermo Fisher Scientific). Approximately 770 evenly spaced measurements of fluorescence against the temperature gradient were recorded and processed by QuantStudio V1.3 (Thermo Fisher Scientific), and then further analyzed by Protein Thermal Shift Software v1.3 (Thermo Fisher Scientific). Thermal shift (Δ*T*_*m*_) was calculated by subtracting the TcPINK1 KD *T*_*m*_ in the presence of a compound to the plate- and column-specific buffer control.

### In vitro inhibitor screen

Inhibitors were purchased from different sources via Cedarlane Laboratories (catalog # in parenthesis). Selleck Chemicals LLC: PRT062607 (S8032), CYC116 (S1171), CYT387 (S2219), URMC-099 (S7343), JNJ-7706621 (S1249), AZD3463 (S7106), VE-822 (S7102), JNK-IN-8 (S4901), staurosporine (S1421), foretinib (S1111); Adooq Bioscience: PRT060318 (A15524-5), TAK-659 (A21885-2). The racemic PRT-core was custom synthesised by ChemSpace (CSCS02808122161). Kinase activity assays were performed by mixing 5 nM of purified TcPINK1 with 100 μM ATP, 100 μM small molecules from 10 mM stock in DMSO, 200 μM MgCl2 in kinase buffer (final 1% DMSO). The reactions were left in a 30°C incubator for 5 minutes for TcPINK1. The reactions were stopped with the addition of Laemmli buffer (2% SDS, 0.1% bromophenol blue (SigmaAldrich), 10% glycerol, 100 mM DTT) final concentration 1X. The entire reaction was loaded onto a 12.5% Tris-Tricine gel (1 M Tris-HCl pH 8.45, 12.5% acrylamide, 0.1% SDS, 0.1% APS, 0.04% TEMED). Protein from a 12.5% polyacrylamide gel were transferred onto an Immuno-blot PVDF membrane (Bio-rad). The membranes were stained with 1% Ponceau and imaged using the ImageQuant LAS 500. The membrane was washed with distilled water and then blocked in 5% BSA diluted in tris buffered saline with 0.1% Tween (TBS-T) for 1 hour. The membrane was briefly rinsed and left to incubate with primary antibody (pUb; Cell Signaling Technology, cat #62802) at a 1:10,000 dilution in TBS-T for two hours. The membrane was further rinsed three times for 10 minutes with TBS-T before incubating it with a 1:2000 dilution of secondary antibody, anti-rabbit, in TBS-T. The membrane was rinsed three times for 10 minutes in TBS followed by the addition of 1 mL ECL solution (BioRad) to the membrane and visualized on the ImageQuant LAS 500.

### NMR spectroscopy

Purified TcPINK1 at 1 μM was combined with 100 μM ^13^C-ATP (Sigma-Aldrich), 100 μM compound of interest, 500 μM MgSO_4_, 5% D_2_O (Sigma-Aldrich), and 5% DMSO in in 300 mM NaCl, 50 mM Tris-HCl pH 8, 2 mM DTT. For the ADP control, knowing that the natural abundance of ^13^C is 1.1%, 1 μM ^15^N-TcPINK1 was combined with 10 mM ADP (Sigma-Aldrich) in the same buffer to re-create a ^13^C signal equivalent to 100 μM ^13^C-ADP. Standard 2D ^1^H-^13^C sensitivity-enhanced HSQC NMR spectra were acquired at 298K every 5 minutes for 1.5 hr on a 600 MHz Bruker Avance spectrometer equipped with a triple resonance (^15^N/^13^C/^1^H) cryoprobe. The spectra were acquired with a carrier frequency of 600.3328216 MHz (4.7 ppm) for F2 (^1^H) and 150.9697038 MHz (110 ppm) for F1 (^13^C); a sweep width of 13.0136 ppm for F2 (^1^H) and 36 ppm for F1 (^13^C); 42 increments and 2 scans. Spectra were processed using TopSpin 4.0.6 (Bruker).

### Autophosphorylation assay and mass spectrometry

Intact protein mass spectrometry was used to monitor transphosphorylation of TcPINK1-D337N by ^15^N-TcPINK1-WT, as described previously (10). ^15^N-TcPINK1 at 5 μM was combined with 5 μM TcPINK1-D337N, 100 μM compound of interest, 1 mM ATP, 2 mM MgSO_4_ in 300 mM NaCl, 50 mM Tris-HCl pH 8, 2 mM DTT with 2% DMSO at 30°C for transphosphorylation reactions with timepoints of 0 s, 30 s, 60 s, and 120 s. Timepoint reactions were stopped by addition of 0.05% TFA and 2% acetonitrile (final concentrations). Reactions in 20 μL were injected on a Dionex C4 Acclaim 1.0/15 mm column followed by a 10 minute 4-50% gradient of ACN in 0.1% formic acid with a flow rate of 40 μL/min. The eluate was analyzed on a Bruker Impact II Q-TOF mass spectrometer equipped with an Apollo II ion funnel ESI source. The multiply charged ions were deconvoluted at 10,000 resolution using the maximum entropy method. The signal intensity of the nonphosphorylated and phosphorylated TcPINK1-D337N peaks were estimated using the DataAnalysis software integration tool and used to calculate the fraction phosphorylated.

### Isothermal titration calorimetry

The isothermal titration calorimetry (ITC) assay was performed using the Microcal ITC 200-1 instrument. The protein sample consisted of TcPINK1 with 4 aromatic mutations for increased solubility (W131A, W142A, Y225A, F401A) prepared in 300 mM NaCl, 50 mM Tris, 2 mM MgCl_2_ and 1 mM TCEP in pH 8.0. Experiment involved 29 injections of AMP-PNP at 1.5 mM into a sample cell containing 350 μL of 100 μM TcPINK1-4arom at 20°C. Data was fitted to a one-site model and background was subtracted using the Origin v7 software.

### In vitro IC_50_ determination (Kinase Glo assay)

The Kinase Glo Max (cat #V6071) kit assay was used to measure the ATP consumed in the kinase assays. Unphosphorylated 1 μM TcPINK1 was incubated with 10 μM ATP for 5 minutes at 30°C in kinase buffer (50 mM Tris, 300 mM NaCl, 3 mM DTT, pH 8.0). Inhibitor (ranging concentration between zero and 100 μM for PRT062607, PRT060318 and TAK659, 400 μM for JNJ-7706621 and 1300 μM of CYC116) and serially diluted in kinase buffer 4-fold 12 times, the 13^th^ sample having no kinase (negative control). The kinase buffer consists of 50 mM Tris-HCl pH 7.5, 300 mM NaCl, 1 mM DTT. 100 μL of kinase at 1 μM was added to 100 μL kinase buffer with inhibitor and left in a 30°C incubator. After the 5-minute mark, 200 μL kinase glo was added to stop the reaction. 100 μL of the sample was then pipetted into 3 wells of an opaque white 96-well plate. The luminescence was measured using the Orion II Microplate Luminometer and obtained using the Simplicity 4.2 program. For each *IC*_*50*_ determination, a corresponding ATP standard curve ranging from zero to 25 μM was generated (Supplemental Fig. S2). The ATP concentration remaining in the reaction was calculated from the linear equation of the ATP standard curve. This number was then divided by the amount of ATP initially present and multiplied by 100 to obtain the percent inhibition. The values were normalized according to the baseline (no inhibitor). The points plotted on a non-linear regression graph in GraphPad Prism 7. The *IC*_*50*_ value and standard deviation were determined in GraphPad.

### In organello assay

Wild type (WT) HeLa cells were treated with 20 μM CCCP for 3 hours to induce the accumulation of PINK1. Cells were then resuspended in mitochondrial isolation buffer (20 mM HEPES, 70 mM sucrose, 220 mM mannitol, pH 7.4, 1 mM EDTA, Complete protease inhibitors) and lysed using nitrogen cavitation for 5 min at 500 psi. Cell debris were removed by centrifugation at 500 g for 5 min, and mitochondria were pelleted by centrifugation at 20,000 g for 20 min. 7.5ug of mitochondria was incubated with 3.75 μg Ub4, 5 mM MgCl2, 0.1 mM ATP and 100 μM of each inhibitor for 15 minutes at 30°C. The reaction was quenched with Laemlli buffer and loaded on a gel for western blotting (described above).

### Cell based assay and IC_50_ determination

WT HeLa cells were transfected with PINK1 for 14h and then treated with a range of PRT concentrations alongside 20 μM of CCCP for 4 h. The cells were then washed and lysed with RIPA buffer (20 mM Hepes, 100 mM NaCl, 0.1% Triton-X, 0.2% SDS, PhosStop, cOmplete inhibitor tablet). Cells were washed with 1 mL PBS twice and lysed with 5 x pellet volume of lysis buffer. The samples were left on ice for 30 minutes then spun at 14,000 rpm for 30 minutes. The supernatant’s concentration was measured with a BCA protein assay kit (J63283.QA). 10 μg of the sample was loaded on a gel for further western blotting (protocol above. Each resulting blot was normalized according to the no inhibitor condition and the highest concentration for the 100% and 0% activity respectively. The data point for the replicates were plotted and the value of the *IC*_*50*_ and its standard error on GraphPad Prism 7.

### Crystallization

PRT060207 was co-crystallized with the TcPINK1 121-570 crystallization construct reported previously (TcPINK1-6arom^121-570^) (8). This construct contains 6 mutations or surface exposed aromatic residues to alanines (W131A, W142A, Y225A, Y378A, F401A and F437A) to enhance solubility and crystallization. CYC116 was co-crystallized with a construct with two additional mutations to TcPINK1-6arom^121-570^: Y144A and I147A. Both crystallization constructs were co-expressed with lambda phosphatase and purified as described above. Crystals of the TcPINK1-6arom^121-570^:PRT060207 were obtained by mixing 0.3 μL of 2.5 mg/ml protein and 100 μM PRT060207 in the presence of 5% DMSO with 0.3 μL 0.1 M HEPES (pH 7.0), 20% PEG 4K and 0.15 M ammonium sulfate, at 22°C using the sitting-drop vapor diffusion method. Crystals of the TcPINK1-6arom^121-570^-Y144A-I147A:CYC116 complex were obtained by mixing 0.8 μL of 5 mg/ml protein and 100 μM CYC116 in the presence of 5% DMSO with 0.8 μL 90 mM HEPES 7.0, 17.5% PEG 4K, 0.13 M Ammonium sulfate, at 16°C using the sitting-drop vapor diffusion method. Both crystals were fished and cryo-protected in mother liquor supplemented with 20% PEG400.

### X-ray structure determination

Diffraction data for both structures were collected on the 24-ID-C beamline at the Advanced Photon Source. 900 images were collected with an oscillation angle of 0.2° at 0.987 Å wavelength. Reflections were processed with *autoPROC* (50), merged, and scaled with *Aimless* (51, 52). The structures were solved by molecular replacement using the structure of apo TcPINK1-6arom^121-570^ (PDB 7MP8) and refined with the *Phenix* suite (53). Models were built with *Coot* (54). Data collection and refinement statistics are given in Table 1. Omit maps were calculated using *Polder* in the Phenix suite. The PRT060207-bound and CYC116-bound structures were deposited to the Protein Data Bank (code 8UCT and 8UDC, respectively).

### Molecular docking

TAK-659 and JNJ-7706621 were docked on the apo structure of TcPINK1^121-570^ (PDB 7MP8) using the software *DiffDock* (55). We used 20 inference steps and generated 40 sample models. The lowest energy model was used for structural analysis.

## Data availability

All data is located in the article. The coordinates and maps for the two structures presented have been deposited in the Protein Data Bank and are publicly available (PDB codes 8UCT and 8UDC). Further information and requests for raw data and materials can be requested from the main contact, Jean-François Trempe.

## Supporting information

This article contains supporting information (Fig. S1-S3).

## Acknowledgments

We thank the staff at the Advanced Photon Source (APS) for help with X-ray data collection. We thank the McGill Pharmacology SPR/MS facility (Mark Hancock) for support in mass spectrometry, and the CRBS for help with the ITC (Kim Munro) and NMR (Tara Sprules).

## Author contributions

Protein purification and crystallization, S.R.; crystallography and structure determination, S.R. S.V.; biochemical and cell-based assays, T.S. and L.T.; thermal shift screening, N.C.; writing & conceptualization, S.R., T.S. and J.-F.T.

## Funding and additional information

This work was supported by a Canada Research Chair (Tier 2) in Structural Pharmacology to J.-F.T., as well as grants from CIHR (153274), Parkinson Canada (2017-1277) and the Michael J. Fox Foundation (12119). S.R. was supported by a studentship from Parkinson Canada, as well as from the Centre de Recherche en Biologie Structurale (Fond de Recherche du Québec – Santé, FRQS). S.V. was supported by an FRQS postdoctoral fellowship and Parkinson Canada Basic Research Fellowship. L.T. was supported by a FRQS scholarship. This research used resources of the Advanced Photon Source, a U.S. Department of Energy (DOE) Office of Science user facility operated for the DOE Office of Science by Argonne National Laboratory under Contract No. DE-AC02-06CH11357.

