## Supplemental Figures for "Identification and structural characterization of small-molecule inhibitors of PINK1"

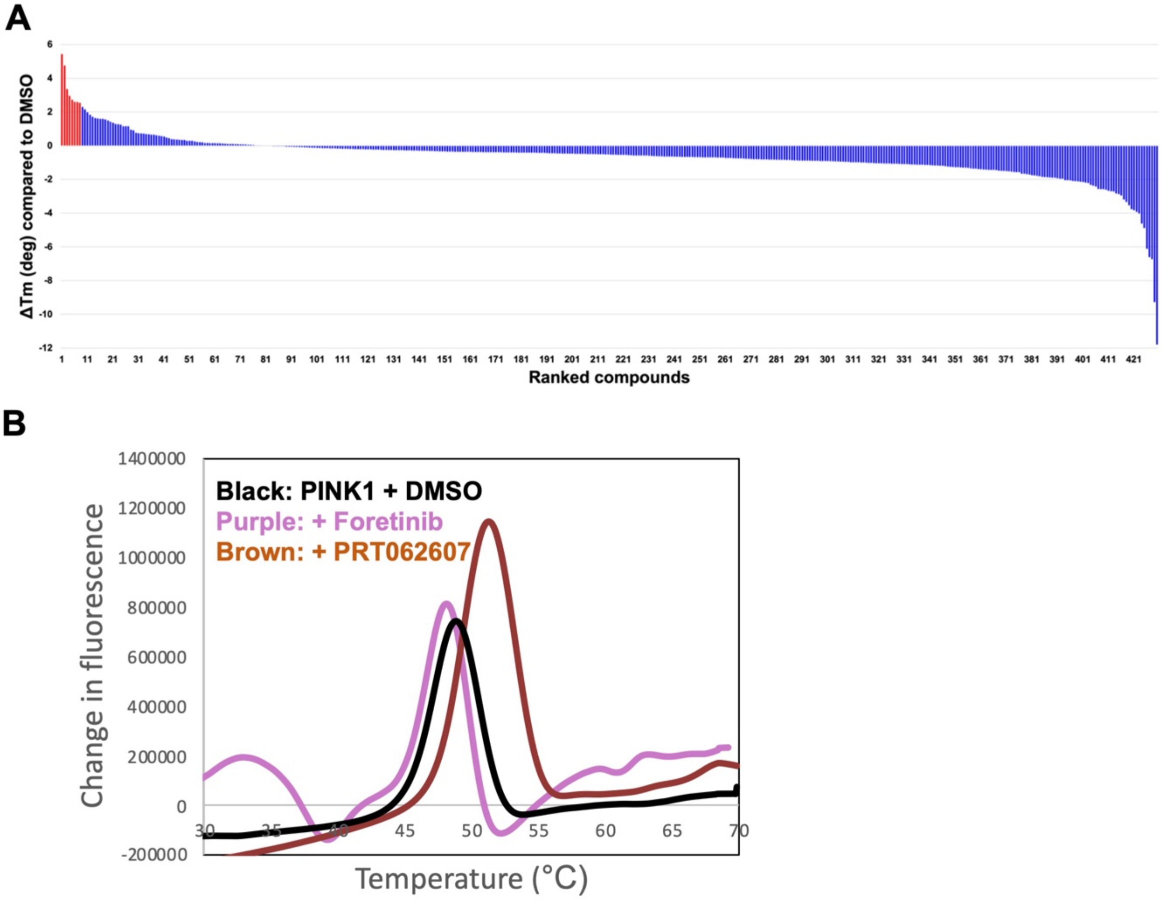


**Supplemental Figure S1. Thermal shift assay screening of TcPINK1 targeting molecules. (A)** Thermal shift (*ΔTm*) values obtained for TcPINK1 incubated with Sypro Orange and 100 µM compound derived from the SelleckChem kinase small molecule library (430 compounds). Compounds were ranked by *ΔTm* value. Top ranked compounds are shown in red. **(B)** Example of raw thermal denaturation data obtained for TcPINK1 incubated with DMSO (baseline), Foretinib (negative control) and PRT062607 (hit, positive *ΔTm*). The first derivative of the change in fluorescence is plotted as a function of the temperature. The Tm corresponds to the peak of the change in fluorescence (inflection point).

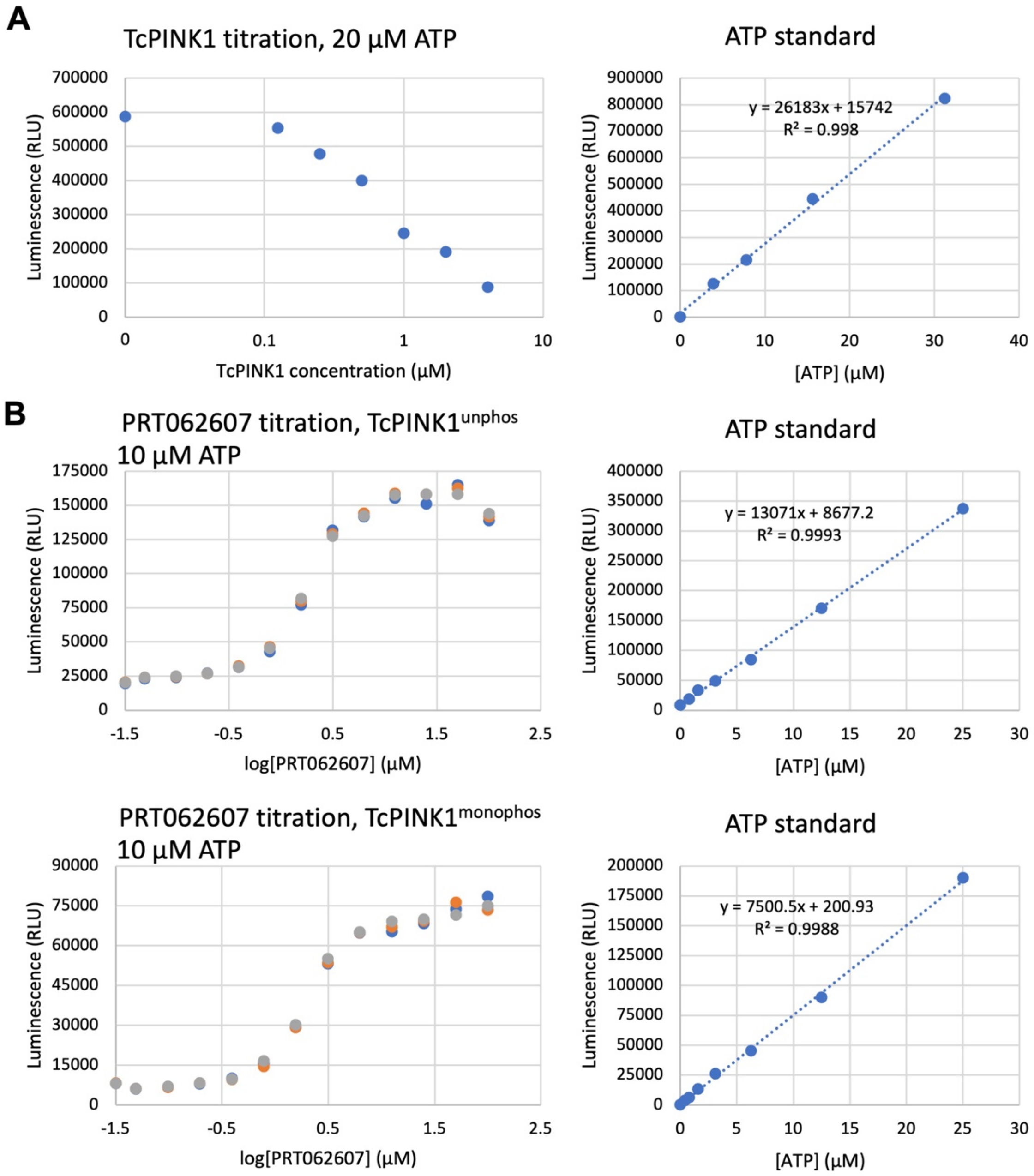


**Supplemental Figure S2. Optimization and analysis of the Kinase Glo assay for *$IC_{50}$* determination. (A)** Luminescence values observed after incubating 20 µM ATP for 5 min in the presence of difference TcPINK1 concentrations (left). The midpoint *$EC_{50}$* is around 1 µM TcPINK1. The ATP standard curve used in this assay is shown to the right and shows linearity for the entire range of luminescence observed. **(B)** Raw luminescence data used for *$IC_{50}$* determination (see Figure 2D). Different concentrations of PRT062607 was incubated with 1 µM $TcPINK1^{121\text{-}570}$ (unphosphorylated or mono-phosphorylated) and 10 µM ATP for 5 min. Reactions were performed in triplicates. The ATP standard curve for each experiment is shown on the right.

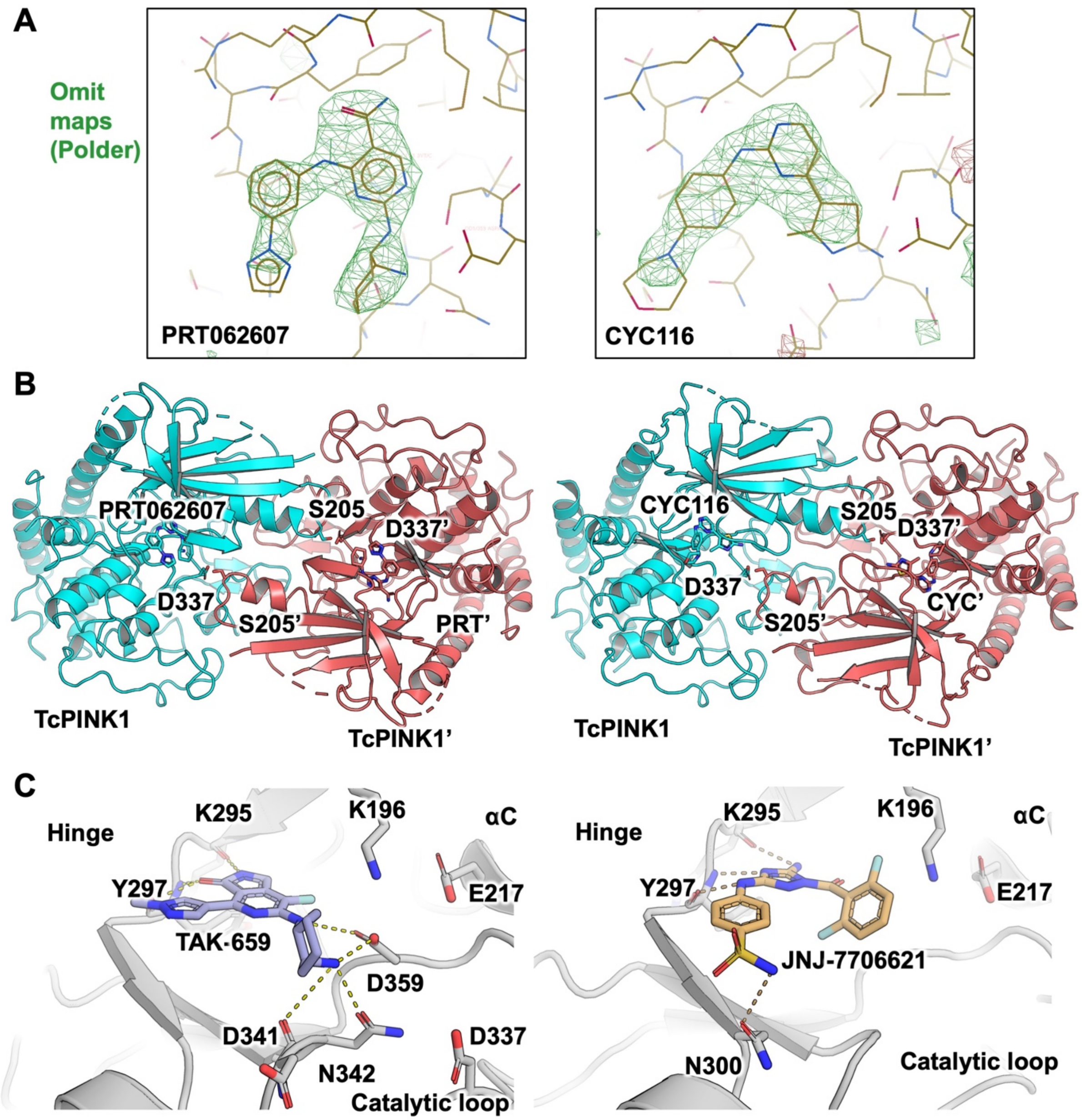


**Supplemental Figure S3. Structural analysis of TcPINK1 binding to different inhibitors. (A)** Polder ligand omit maps (3.5 σ) calculated from the crystal structures TcPINK$^{121\text{-}570}$ bound to PRT062607 (left) and CYC116 (right). **(B)** Crystallographic symmetry reveals face-to-face trans autophosphorylation complex in both PRT060207 and CYC116-bound TcPINK1, as observed previously for the apo and AMP-PN bound structures (PDB: 7MP8 and 7MP9). **(C)** Docking models of TcPINK1 bound to TAK-659 (left) or JNJ-7706621 (right), showing their interactions with the hinge, A-loop and catalytic loop. The apo structure of TcPINK1$^{121\text{-}570}$ (PDB 7MP8) was used for docking the small molecules using the software *DiffDock*.
